# A Bayesian Approach to Beam-Induced Motion Correction in Cryo-EM Single-Particle Analysis

**DOI:** 10.1101/384537

**Authors:** Jasenko Zivanov, Takanori Nakane, Sjors H. W. Scheres

## Abstract

We present a new method to estimate the trajectories of particle motion and the amount of cumulative beam damage in electron cryo-microscopy (cryo-EM) single particle analysis. We model the motion within the sample through the use of Gaussian Process regression. This allows us to associate with each hypothetical set of particle trajectories a prior likelihood that favours spatially and temporally smooth motion without imposing hard constraints. This formulation enables us to express the a-posteriori likelihood of a set of particle trajectories as a product of that prior likelihood and an observation likelihood given by the data, and to then maximise this a-posteriori likelihood. Since our smoothness prior requires three parameters that describe the statistics of the observed motion, we also propose an efficient stochastic method to estimate those parameters. Finally, we propose a practical means of estimating the average amount of cumulative radiation damage as a function of radiation dose and spatial frequency, and a robust method of fitting relative B-factors to it. We evaluate our method on three publicly available datasets, and illustrate its usefulness by comparison with state-of-the-art methods and previously published results. The new method has been implemented as Bayesian polishing in RELION-3, where it replaces the existing particle polishing method, as it outperforms the latter in all tests conducted.

## 1 Introduction

Recent advances in electron detector technology have allowed cryo-EM single-particle analysis to uncover the structures of many biological macromolecules to resolutions sufficient for *de novo* atomic modelling. The primary impediment to high-resolution reconstruction is radiation damage that is inflicted on the molecules when they are exposed to an electron beam. This requires low-dose imaging, and hence reconstructions from very noisy images. In addition, exposure to the electron beam leads to motion in the sample, which destroys information, particularly in the high spatial frequencies.

Because new detectors allow capturing multi-frame movies during exposure of the sample, it is possible to estimate and correct for beam-induced motion. This requires sufficient signal in the individual movie frames, which is challenging as each frame only contains a fraction of the total electron dose, resulting in even lower signal-to-noise ratios. The earliest approaches to beam-induced motion correction were performed in *Frealign* [4, 5] and relion [2], and estimated particle positions and orientations independently in each movie frame and for each particle. Both programs averaged the signal over multiple adjacent frames to boost the low signal-to-noise ratios. Still, these approaches were only applicable to relatively large (> 1 MDa in molecular weight) particles (i.e. molecules or molecular complexes). These early studies revealed correlations between the direction and extent of motion of particles that are in close proximity to each other. In this paper, we will refer to this property as the *spatial smoothness* of motion.

The approach in [2] was subsequently extended to cover smaller molecules. This was possible by (1) still performing template matching on averages over multiple adjacent frames; (2) fitting a linear path of constant velocity through the unreliably detected positions; and (3) averaging those constant velocity vectors over local areas of the micrograph. This means that consistency with the observations and the (absolute) temporal and (partial) spatial smoothness of the trajectories were imposed one after the other. That algorithm, together with a radiation-dose weighting scheme described below, was termed particle polishing [21], and was implemented as the method of choice for beam-induced motion correction in the Relion package [20].

In the meantime, a second class of motion estimation algorithms have been developed that do not rely on the availability of a three-dimensional reference structure, and which, therefore, can be applied much earlier in the image-processing workflow. Instead of comparing individual particles with their reference projections, these algorithms estimate the motion entirely from the frame sequence itself, by cross-correlating individual movie frames or regions within them. This tends to work best for relatively low spatial frequencies, which can be challenging for movies that are captured close to focus, where the low frequencies are the most attenuated by the contrast transfer function.

Two of the early reference-free methods, *MotionCorr* [10] and *Unblur* [7], relaxed the spatial smoothness assumption, allowing for non-linear trajectories. While *MotionCorr* allowed completely free motion over time, it required discrete regions of the image to move as rigid blocks. *Unblur* imposed a certain amount of temporal smoothness on the motion and it required the entire image to move as a rigid block. The method by Abrishami et al. [1] was based on an iterative version of the Lucas-Kanade optical flow algorithm [12] and it abandoned the idea of rigid regions in favour of a model that allows for spatially smooth deformations of the image. Later, a more robust noise model was proposed in *Zorro* [14], which required uniform movement of the entire micrograph, and a variant of it, *SubZorro*, that worked on rigid regions.

An early method to formulate motion estimation as a minimisation of a cost function in order to simultaneously satisfy consistency with the observations and temporal smoothness was *alignparts-lmbfgs* [19]. It estimated the motion of each particle separately, so spatial smoothness of the motion was enforced only after the fact, by forming local averages over trajectories of neighbouring particles. Although *alignparts-lmbfgs* works on individual particles, the program does not use reference projections, but minimises a weighted phase difference between the Fourier components of individual movie frames of boxed out particles.

A reference-free method that is very popular today is *MotionCor2* [25]. This program enforces neither spatial nor temporal smoothness absolutely. Instead of working on individual particles, it splits the micrograph into tiles, and it fits the motion of each tile to a global polynomial function of time and space. This is done by picking independent, most likely positions of each block and then fitting the coefficients of the polynomial to those discrete positions. We will compare our new method to *MotionCor2* in the results section.

Unlike particle motion, radiation damage cannot be corrected for explicitly. Nevertheless, the deleterious effects of radiation damage on the reconstruction can be reduced by down-weighting the contribution of the higher spatial frequencies in the later movie frames. This is because radiation damage affects the signal at high spatial frequencies faster than the signal at low spatial frequencies [8]. For this reason, it was proposed to discard the later movie frames for high-resolution reconstruction [10]. The particle polishing program in relion [21] would then extend this to a continuous radiation-damage weighting scheme. This approach used a relative B-factor model (based on the temperature factors that are commonly used in X-ray crystallography) to describe the signal fall-off with resolution. Later, Grant and Grigorieff derived a more precise exponential damage model from a reconstruction of a rotavirus capsid, and proposed this as a general model for radiation damage in biological cryo-EM samples [7]. The latter is currently in use in many programs.

In this paper, we describe a new method, which we have termed Bayesian polishing, and which has been implemented in the relion package. This method still uses the original B-factor model for the relative weighting of different spatial frequencies in different movie frames, although we do propose a new method to estimate the B-factors. We chose the B-factor model because, as opposed to the exponential model by Grant and Grigorieff, it allows us to model both radiation damage and any residual motion that is not corrected for. However, as the B-factors can only be determined once the motion has been estimated, we do use the exponential model during the initial motion estimation step.

The two main disadvantages of the motion estimation process in the original particle polishing algorithm in relion that prompted these developments were the absolute temporal smoothness assumption and the feed-forward nature of the fitting process: a linear path that best fits the estimated noisy positions might not be the linear path that leads to the greatest overall consistency with the observed data. In other words, the per-frame maxima are picked prematurely. This is illustrated in Fig. 1. The same is also true for the spatially smooth velocity field that results from the averaging of multiple such linear trajectories. The motion estimation method that we propose in this paper overcomes both of these disadvantages.

**Figure 1:**
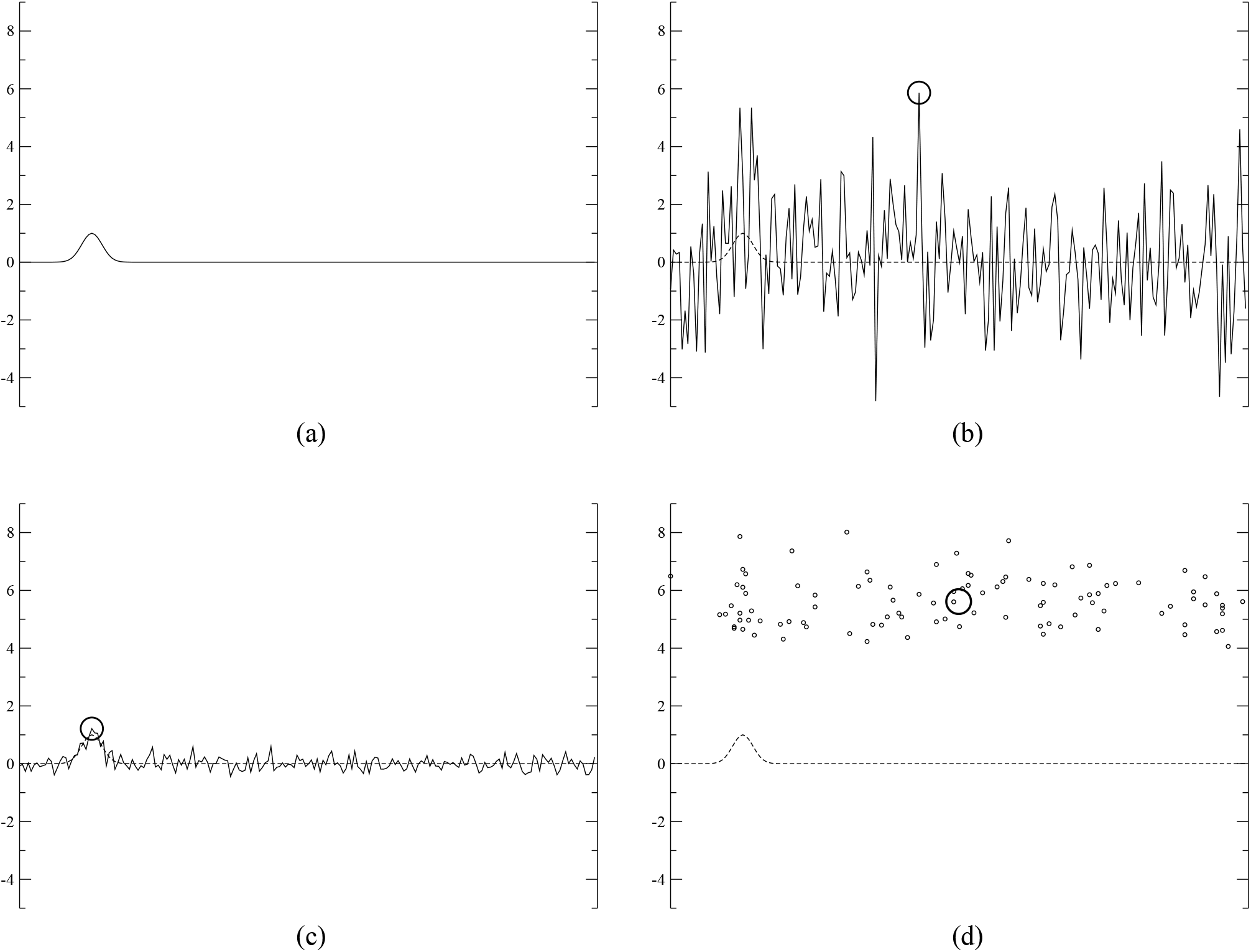
*A simulated example illustrating the issue of premature maximum picking. **(a)**: A Ga,ussia,n representing the cross-correlation between the reference and the observation of a particle. **(b)**: That cross-correlation distorted by Gaussian white noise of realistic intensity for the cross-correlation between a noise-free reference and one observed frame of one single particle (σ* = 2; *the circle indicates the maximum). **(c)**: The average over* 100 *such noisy functions and its maximum. **(d)**: The maxima of those* 100 *noisy functions (small circles) and their average (big circle). Note how the average of the noisy maxima (d) is much further from the true maximum than the maximum of the average (c). Our proposed method avoids picking individual maxima of noisy functions, and it instead aims to maximise the cross-correlations of all particles and the prior smoothness assumptions simultaneously*.

## 2 Materials and Methods

In the following, we will discuss the different components of our proposed Bayesian polishing approach. We will begin by describing the motion model and the motion estimation process in 2.1. After that, we will explain how parameters for our prior, i.e. for the statistics of motion, are determined in 2.2. Although those have to be known in order to estimate the most likely motion, we chose to describe their determination afterwards, since its understanding requires knowledge of the actual motion model. We then describe the process of measuring the relative B-factors and recombining the frames in 2.3, and we conclude this section with a description of our evaluation process in 2.5.

### 2.1 Motion Estimation

#### 2.1.1 Outline

The central idea behind our motion estimation consists in finding a set of particle trajectories in each micrograph that maximise the a-posteriori probability given the observations. Note that we assume a reference map, the viewing angles and defoci of the particles and the parameters of the microscope to be known by this point.

Formally, we express the particle trajectories as a set of positions 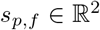, for each particle *p* ∈ {1..*P*} and frame *f* ∈ {1..*F*}. The corresponding per-frame particle displacements are denoted by *v_p,f_* = *s*_*p,f* + 1_ − *s_p,f_* for *f* ∈ {1..*F* − 1}. We will refer to *v_p,f_* as per-frame velocities in the following, since they are equal to the mean velocities between the two frames if time is measured in units of frames.

Let *s* = {*s_p, f_*|*p* ∈ {1..*P*}, *f* ∈ {1..*F*}} denote the set of all particle trajectories in a micrograph. The a-posteriori probability *P*_AP_(*s*|`obs`) of those trajectories given the observations `obs` is then given by Bayes’ law:

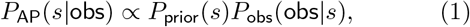

where the term *P*_prior_(*s*) describes the prior probability of that set of trajectories and is described by the statistics of motion, while *P*_obs_(`obs`|*s*) describes the probability of making the observations *obs* given those trajectories.

We will first describe our motion model that gives rise to *P*_prior_(*s*) in 2.1.2, and then the observation model that defines *P*_obs_(`obs`|*s*) in 2.1.3.

#### 2.1.2 The Motion Model

We model particle motion using Gaussian-Process (GP) regression. GPs have been in use by the machine learning community for decades [17], and they have found applications in the fields of computer vision [13], computer graphics [23] and robotics [16].

Formally, a GP is defined as a distribution over the space of functions *f*(*x*) such that for every finite selection of *x_i_*, the corresponding *f*(*x_i_*) follow a multivariate normal distribution. A GP can therefore be thought of as an extension of the concept of a multivatiate normal distribution to cover the (infinitely-dimensional) Hilbert space of functions. Although the term process suggests *x* being a one-dimensional time variable, a GP can in fact be defined over any domain. In our case, we use the particle positions in the micrograph (i.e. a 2D plane) as that domain, while the function *f*(*x*) will be used to describe the velocity vectors of particles.

In its most general form, a GP is defined by a mean *μ*(*x*) and a covariance function *C*(*x*_1_, *x*_2_). In our specific case, we will assume the mean velocity to be zero, and we will work with *homogeneous* GPs, where the covariance between two points *x*_1_ and *x*_2_ depends only on their distance *d* = |*x*_2_ − *x*_1_|. We will use the GP to enforce spatial smoothness of the motion vectors. This means that the covariance *C*(*d*) between two velocity vectors will be greater for particles that are closer together.

Specifically, the covariance between the velocities of two particles *p* and *q* is modelled by the exponential kernel:

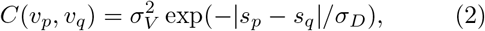

where *σ_V_* describes the expected amount of motion, while *σ_D_* describes its spatial correlation length. Since the overall beam-induced motion of the particles is generally far smaller than their mutual distance (a few Å vs. hundreds of Å), we chose to compute the covariance based on the initial particle positions alone – this is why the subscript *f* is missing in Eq. 2.

We can write the covariances of all particles *C*(*v_p_*, *v_q_*) into a *P × P* covariance matrix Σ*_V_*, which then describes the per-frame multivariate normal distribution of all velocity vectors *v_p,f_*. As is commonly done in GP regression, we perform a singular-value decomposition on Σ*_V_* to obtain a more practical parametrization for our problem:

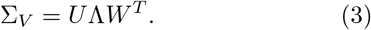

This allows us to define a set of basis vectors 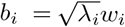, where 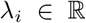 is the *i^th^* singular value and 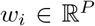 is its associated singular vector (i.e. column of *W* or row of *W^T^*). For each frame, the *x* and *y* component of the velocity vectors *v_p_* of all particles *p* can now be expressed as linear combinations of *b_i_* with a set of *P* coefficients *c_i_*:

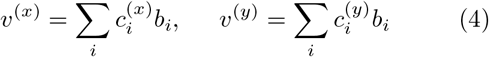

In this parametrization, the per-frame joint likelihood of that set of velocities has a particularly simple form:

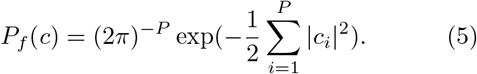

For this reason, we use *F* − 1 sets of coefficients *c_i,f_* as the unknowns in our problem. Since the *c_i_* only describe the velocities, they only determine the positions *s_p,f_* up to a per-particle offset. The complete set of unknowns for a micrograph therefore also has to include the initial positions *s*_*p*,0_. The initial positions have no effect on the prior probability, however.

Formally, for 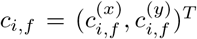, the positions are then given as a function of all *c_i,f_* by:

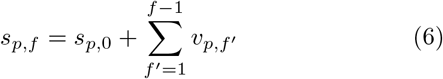

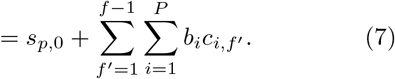

So far, we have only modelled the spatial smoothness of the motion. To impose temporal smoothness, we define the complete prior probability as

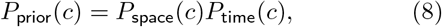

with

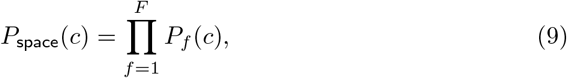

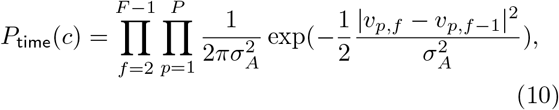

where *σ_A_* is the third and final motion parameter that describes the average acceleration of a particle during a frame, i.e. the std. deviation of the change in velocity between two consecutive frames.

The temporal smoothness term *P*_time_ corresponds to the one proposed by Rubinstein and Brubaker [19] for individual particles. From the orthogonality of the basis *b_i_*, it follows that in our parametrization,

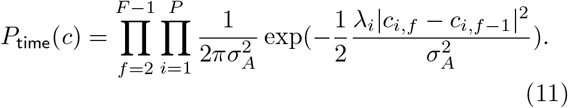

#### 2.1.3 The Observation Model

In the following, we will derive the observation likelihood *P*_obs_(`obs`|*x*). Since we assume a 3D reference map, the viewing angles and the microscope parameters to be known, we can predict the appearance of a particle using the reference map [20]. This is done by integrating the reference map along the viewing direction, which can be accomplished efficiently by extracting a central slice in Fourier space, and then convolving the resulting image with the known contrast transfer function (CTF).

To maintain the nomenclature from previous Relion papers, we denote pixel 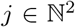 of frame *f* of the experimental image of particle *p* by *X_p,f_*(*j*) and the same pixel in the prediction by *V_p,f_*(*j*). The spectral noise power is measured from all *X* in a micrograph, and both *X* and *V* are filtered in order to whiten the image noise (i.e. decorrelate the noise between the pixels) and to scale it to unit variance. In addition, we use the exponential damage model [7] to suppress the high frequencies in the later frames in *V*.

By assuming that the noise in the pixels is Gaussian and independent, it follows that

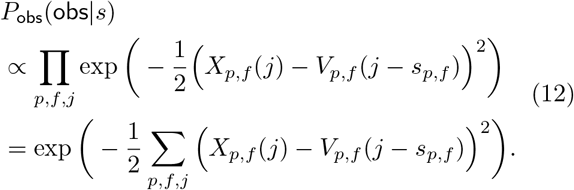

Since the prediction *V* is zero outside of the molecule, the image area over which this sum is evaluated only influences the scale of *P*_obs_, but not its shape. In practice, we cut out a square from the micrograph that contains the molecule (including a certain amount of padding around it to account for its motion) and evaluate *P*_obs_ on that square.

In order to evaluate *P*_obs_(`obs`|*s*) efficiently for different hypothetical particle positions *s*, we use the following identity:

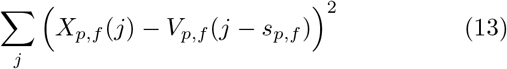

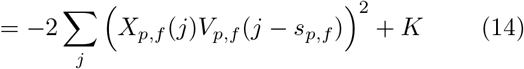

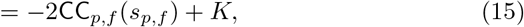

where `CC`_*p,f*_ denotes the cross-correlation between *X_p,f_* and *V_p,f_*, which is computed for a Cartesian grid of integral *s* simultaneously via a convolution in Fourier space. The constant offset *K* merely scales the resulting probability *P*_obs_, so it does not alter the location of the maximum of *P_AP_* = *P*_prior_*P*_obs_. We can thus define

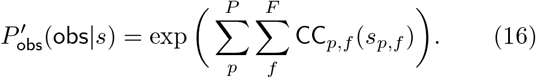

To determine the values of `CC`_*p,f*_ at fractional coordinates, we apply cubic interpolation. This ensures a continuous gradient.

#### 2.1.4 Optimisation

To avoid numerical difficulties, we maximise *P*_AP_(*s*|`obs`) by instead minimizing its doubled negative log, *E*_AP_ = −2log(*P*_AP_). The doubling serves to simplify the terms. All the products in *P*_AP_ become sums in *E*_AP_, yielding:

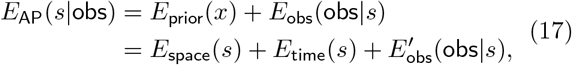

where the terms *E*_pace_, *E*_time_ and *E*′_obs_ are defined analougously. Inserting the terms defined in 2.1.2 and 2.1.3 yields:

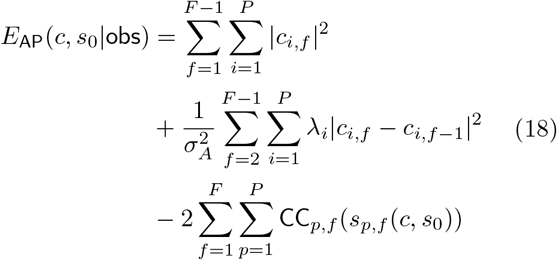

The expression in Eq. 18 is differentiated with respect to the coefficients *c_i,f_* and initial positions *s*_*p*,0_ for all *i* and *f*, and the combination that minimises *E*_AP_(*c*, *s*_0_|`obs`) is determined using the L-BFGS algorithm [11]. In order to avoid overfitting, all particles are aligned against a reference computed from their own independently refined half-set [22].

### 2.2 Parameter Estimation for the Statistics of Motion

The estimation procedure described in 2.1 requires three parameters (*σ_V_*, *σ_D_* and *σ_A_*) for the prior that encapsulate the statistics of particle motion. Since the precise positions of the particles can never be observed directly, measuring those statistics requires performing a process of hyper-optimization, i.e. optimizing motion parameters that produce the best motion estimates. This renders the entire approach an empirical Bayesian one. The simplest solution would be to perform a complete motion estimation for each hypothetical triplet of motion parameters. As the motion estimation usually takes multiple hours on a non-trivial dataset, this would become prohibitive for a 3D-grid of parameters.

Instead, we estimate the optimal parameters using the following iterative procedure. First, we select a representative random subset 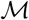 of micrographs that contain at least a pre-defined minimal number of particles (25,000 in our experiments). Then, we perform the following three steps iteratively.

1. Choose a hypothetical parameter triplet *σ_V_*, *σ_D_* and *σ_A_*.
2. Align all micrographs in 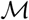 using those parameters.
3. Evaluate the parameters.

The iterations are performed using the Nelder-Mead uphill simplex algorithm [15] which does not rely on the function over which it optimises being differentiable.

In order to evaluate a parameter triplet, we perform the alignment only on a limited range of spatial frequencies (the *alignment circle*, 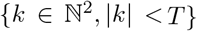). The remainder of frequencies, the *evaluation ring* {|*k*| > *T*}, is used to evaluate that alignment. To avoid overfitting, i.e. to retain a strict separation of the two half-sets, we perform the alignment against a reference obtained from the same half-set to which the respective particle belongs. For the evaluation, we use a reference obtained from the opposite half-set to avoid the particle “finding itself’ [7] in the reference. Note that the latter does not incur any risk of overfitting, since the alignment is already known by the time it is evaluated, and the small number of parameters (i.e. three values) leaves no room for overfitting.

The partition of frequency space into an alignment circle and an evaluation ring is necessary: if the alignment and the evaluation were to be performed on the same frequencies *k*, then a weaker prior would always produce a greater correlation than a stronger one. Note that this would happen in spite of splitting of the particles into independent half-sets, because an insufficiently regularised alignment will align the noise *in the images* with the signal in the reference, while the two references share the same signal in the frequency range in which they are meaningful.

The evaluation itself is performed by measuring what we propose to call the thick-cylinder correlation 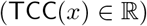 between the aligned images and the reference:

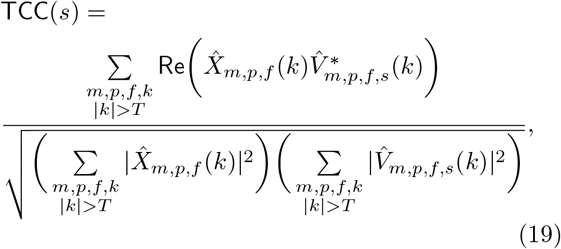

where 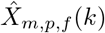 and 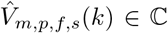 are the Fourier-space amplitudes of frequency *k* of the observed image and the prediction, respectively. The indices denote frame *f* of particle *p* in micrograph 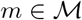. The prediction 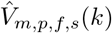 has been shifted according to the estimated *s_m,p,f_*, i.e. 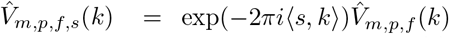. The asterisk indicates complex conjugation and 〈,〉 a 2D scalar product.

### 2.3 Damage Weighting

Once the frames of a movie have been aligned, we compute a filtered average over them that aims to maximise the signal-to-noise ratio in each frequency. In the original particle polishing method [21], the proposed image recombination approach was based on relative B-factors. We use that same approach here, but we propose a more practical and more robust means of estimating the relative B-factors.

The original technique required computing two full 3D reconstructions from particle images of every frame, one for each independently refined half-set. In a typical dataset comprising 40 frames, this would amount to computing 80 individual reconstructions, which requires days of CPU time. The two corresponding reconstructions would then be used to determine the Fourier Shell Correlation (FSC) in order to estimate the spectral signal-to-noise ratio (SSNR) of the 3D reconstruction.

Our new method is more practical in that it avoids the computation of those 3D reconstructions. Instead, we directly measure the correlation between the aligned frames and the reference as soon as the particles in a movie have been aligned. This is done by evaluating what we have termed the Fourier-cylinder correlation `FCC`(*f*, *κ*) for each frame index *f* and Fourier shell *κ*. This amounts to correlating the set of Fourier rings of radius *κ* against the reference for all particles simultaneously, hence the term Fourier cylinder.

Formally, the `FCC` is defined as follows:

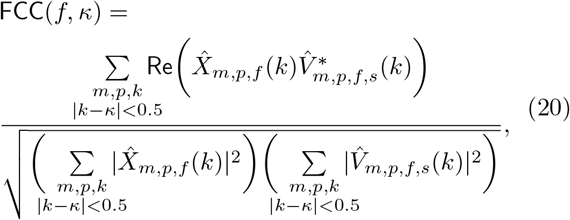

for *k* and *κ* given in pixels. It can be evaluated by iterating over the dataset only once, updating the three sums in Eq. 20 for each particle in each micrograph.

The FCC allows us to estimate the SSNR of the aligned images themselves, not of the 3D reconstructions. The fact that those SSNR values are different is of no concern, as we are only interested in their relative change as a function of frame index *f*. Since the value of each voxel of a 3D reconstruction is an average over the pixels from many images, the relative change in the SNR of that voxel over time is the same as for the corresponding pixels.

Once the `FCC` has been determined, we proceed to fit the relative B-factors. This is done by finding a *B_f_* and 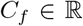 for each frame *f* and a 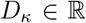 for each frequency ring *κ* that minimise:

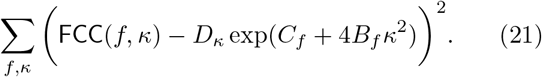

Here, the coefficients *D_κ_* are nuisance parameters that encapsulate the amount of signal in the reference in each frequency band *κ*. This allows the *B_f_* and *C_f_* factors to only express the variation in signal over the frame index *f*. The *D_κ_* are higher for frequencies that are more prominent in the structure (such as those of alpha-helices) and they are zero beyond the resolution of the current reference map. In the previous particle polishing formulation, the D_κ_ correspond to a Gaussian over *κ* given by the average B-factor. The coefficients *B_f_* and *C_f_* maintain the same meaning as in the previous formulation, i.e. the change in high-frequency information and overall contrast over time, respectively. An illustration of the model is shown in Fig. 2.

**Figure 2:**
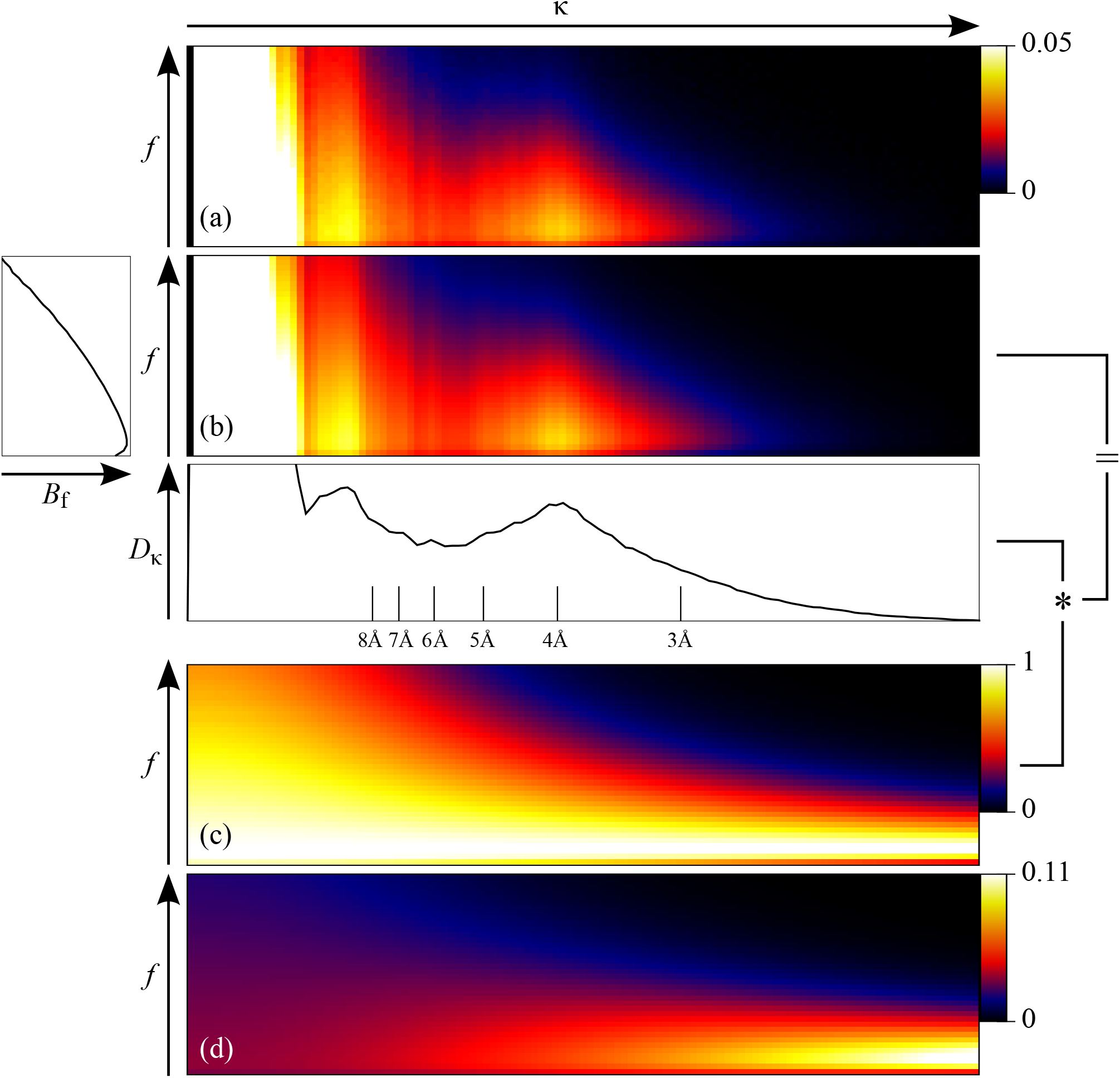
*An illustration of FCC-based B-factor fitting using the *β*-galactosidase dataset as an example (best viewed in colour)*. **(a)***: The FCC computed using Eq. 20 as a function of spatial frequency κ and frame f*. **(b)***: A fit of B_f_, C_f_ and D_κ_ according to Eq. 21 with plots of B_f_ and D_κ_ shown in relation*. **(c)***: The same fit with all D_κ_ set to 1 (i.e. the numerator of Eq.22)*. **(d)***: The normalized weights w_κ,j_ as given by Eq. 22*.

The factors *B_f_*, *C_f_* and *D_κ_* are estimated iteratively, by first finding the optimal *D_κ_* for each *κ* given the current *B_f_* and *C_f_*, and then the optimal *B_f_* and *C_f_* given the current *D_κ_*. The optimal *D_κ_* can be determined linearly, while the *B_f_* and *C_f_* are found through a recursive 1D-search over *B_f_* – the optimal *C_f_* for a given *B_f_* can also be determined linearly. In our implementation, the entire procedure is run for 5 iterations, and it typically takes less than a second to complete.

The final weight of each Fourier-space pixel is then given by:

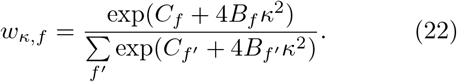

### 2.4 Implementation

The motion estimation algorithm has been implemented using MPI, allowing it to align multiple micrographs in parallel on different computers. The processes that are run on each of those computers are further parallelised using OpenMP, which allows the user to exploit all of the available CPU cores on all of the available computers at the same time. Although it is also possible to align multiple micrographs on the same computer simultaneously by running multiple MPI processes there, we discourage this, since it requires each of those processes to maintain its own data in memory. If the multiple CPU cores of the same computer are instead allowed to cooperate in aligning the same micrograph, then the memory is only taken up once.

The memory footprint of the motion estimation algorithm consists primarily of the two 3D reference maps (one for each independently refined half-set) and the pixels of the micrograph that is currently being processed. In most cases, this requires approximately 20 Gb of memory for each MPI process.

Due to its iterative nature, the parameter hyper-optimisation algorithm does not allow for MPI parallelisation. Furthermore, in order to avoid loading the subset of micrographs from disk in each iteration, all the necessary data are stored in memory. For this reason, the memory footprint of the parameter hyper-optimisation algorithm could exceed 60 Gb for the 25,000 particles used in our experiments. Although a smaller number of particles does reduce this footprint, it also renders the estimated optimal parameters less accurate.

Finally, in order to save disk space, the entire motion estimation pipeline supports micrographs stored as compressed TIFF images. Such images contain the integral numbers of counted electrons for each pixel, which enables very efficient compression: usually, by a factor of about 30. Due to the integral pixel values, an external gain reference has to be provided if such TIFF images are being used.

### 2.5 Experimental Design

We evaluated Bayesian polishing on three publicly available datasets that cover a range of particle sizes: the *P. falciparum* cytoplasmic ribosome (EMPIAR 10028), *E. coli β*-galactosidase (EMPIAR 10061) and human *γ*-secretase (EMPIAR 10194). For all three cases, our group has previously published structures calculated using the original particle polishing approach [24, 9, 3]. We used the same particles and masks for both polishing and the final high-resolution refinement as were used in those papers. Further information on these datasets is shown in Table 1.

**Table 1:**
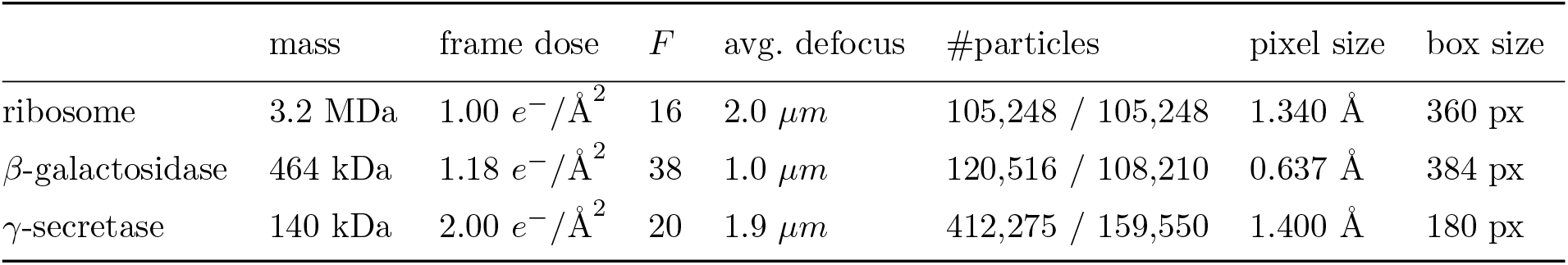
*Properties of the three datasets. The two entries in the particle number column refer to the number used during motion estimation and refinement, respectively*.

| | mass | frame dose | $F$ | avg. defocus | #particles | pixel size | box size |
| --- | --- | --- | --- | --- | --- | --- | --- |
| ribosome | 3.2 MDa | $1.00 \text{ e}^-/\text{\AA}^2$ | 16 | $2.0 \text{ }\mu\text{m}$ | 105,248 / 105,248 | $1.340 \text{ \AA}$ | 360 px |
| β-galactosidase | 464 kDa | $1.18 \text{ e}^-/\text{\AA}^2$ | 38 | $1.0 \text{ }\mu\text{m}$ | 120,516 / 108,210 | $0.637 \text{ \AA}$ | 384 px |
| γ-secretase | 140 kDa | $2.00 \text{ e}^-/\text{\AA}^2$ | 20 | $1.9 \text{ }\mu\text{m}$ | 412,275 / 159,550 | $1.400 \text{ \AA}$ | 180 px |

The experiments were set up as follows. First, the input movies were aligned and dose-filtered using *MotionCor2* [25]. From these aligned micrographs, particles were extracted and an initial reference reconstruction was computed using relion’s 3D auto-refinement procedure [20]. With this reference map, the three parameters that describe the statistics of motion (*σ_V_*, *σ_D_* and *σ_A_*) were determined for each dataset, and the Bayesian polishing algorithm was run on the original, unaligned micrographs. One set of B-factors were estimated for an entire dataset, assigning one B-factor value to each frame index. Using these, a set of motion-corrected and B-factor-weighted particle images were computed, called *shiny* particles in relion, which were then used for a second round of 3D auto-refinement to produce a final map.

Since the official UCSF implementation of *MotionCor2* does not output motion that can be easily interpolated at the positions of the individual particles, we have written our own version of *MotionCor2*. The two implementations are not completely identical. Specifically, our version lacks the fallback mechanism of considering larger tiles if the signal in a tile is insufficient, and it only estimates one set of polynomial coefficients for the entire frame range, while the UCSF implementation always estimates two. In Sec. 3.4, we will show direct comparisons of the FSCs resulting from the two versions to confirm that they give similar resolutions of the final reconstructions.

The particle trajectories for Bayesian polishing were initialised with the motion estimated by our version of *MotionCor2*. This initialisation does not appear to be strictly necessary, however, since in most cases the Bayesian polishing algorithm converged to the same optima if initialised with an unregularised global trajectory. As an example, 90% of the final *β*-galactosidase particle positions showed a difference of less than 10^-4^ pixels as a result of initialisation.

The resulting maps were compared with ones obtained from both versions of *MotionCor2* and with the previously published results. Since the resolution of the resulting maps is influenced by many different factors beyond particle motion, we assume that the estimated relative B-factors reflect the efficacy of motion estimation more reliably than the resolution alone. For that reason, we have also compared the estimated B-factors to those obtained from our version of *MotionCor2* and to the previously published ones. A B-factor comparsion to the UCSF version of *MotionCor2* is not possible, since the particle trajectories are not readily available.

## 3 Results and Discussion

### 3.1 Motion Parameters

The motion parameters were estimated as described in Sec. 2.2. The results are shown in Table 2. We used 25,000 randomly selected particles to estimate the parameters. Performing these calculations multiple times showed that the random subset of micrographs that was used to select the 25,000 particles did affect the outcome of the actual values. Specifically, subsets containing micrographs that exhibited a large amount of stage drift would produce a simultaneous increase in the values of *σ_V_* and *σ_D_*, i.e. stronger and spatially smoother motion. Nevertheless, the choice among different such parameter triplets did not have a measurable impact on the resolution of the resulting reconstructions (results not shown). We assume that stage drift is also the most important reason behind the difference in parameter values among the three datasets, though other reasons might include the size of the molecule and the thickness of the ice.

**Table 2:** *Optimal parameter values used for motion estimation. The values of σv and σ_A_ are normalised by fractional dose (which is given in e^−^/Å)*.

| | $\sigma_V [\text{\AA}/(e^-/\text{\AA}^2)]$ | $\sigma_D [\text{\AA}]$ | $\sigma_A [\text{\AA}/(e^-/\text{\AA}^2)]$ |
| --- | --- | --- | --- |
| ribosome | 1.17 | 28,650 | 1.6 |
| β-galactosidase | 0.66 | 3,300 | 1.5 |
| γ-secretase | 0.57 | 10,710 | 3.0 |

### 3.2 Motion

Using the motion parameters from Table 2, we estimated the motion trajectories for all particles in the three data sets. These calculations took 128 CPU-hours for the ribosome and 778 CPU-hours for *β*-galactosidase on 3.0 GHz Intel Xeon cores, and 1,464 CPU-hours for *γ*-secretase on 2.9 GHz Intel Xeon cores. This is comparable to the computational cost of the existing movie refinement implementation in RELION. Examples of trajectories estimated by Bayesian polishing and our implementation of *MotionCor2* are shown in Fig. 3. A qualitative comparison suggests that they describe the same motion, although they differ in the details. The difference is the most pronounced for *β*-galactosidase, where the motion statistics correspond to very incoherent motion (i.e. a low *σ_D_*). In addition, the trajectories from Bayesian polishing are smoother than the trajectories from *MotionCor2*. This is due to the fact that the global component of the motion is not regularised in *MotionCor2*. The latter has probably no real impact on the resolution of the reconstruction, since the irregularities are far smaller than one pixel. However, quantitative statements about the quality of motion estimation can only be made once a full reconstruction has been computed. This will be done in Sec. 3.4.

**Figure 3:**
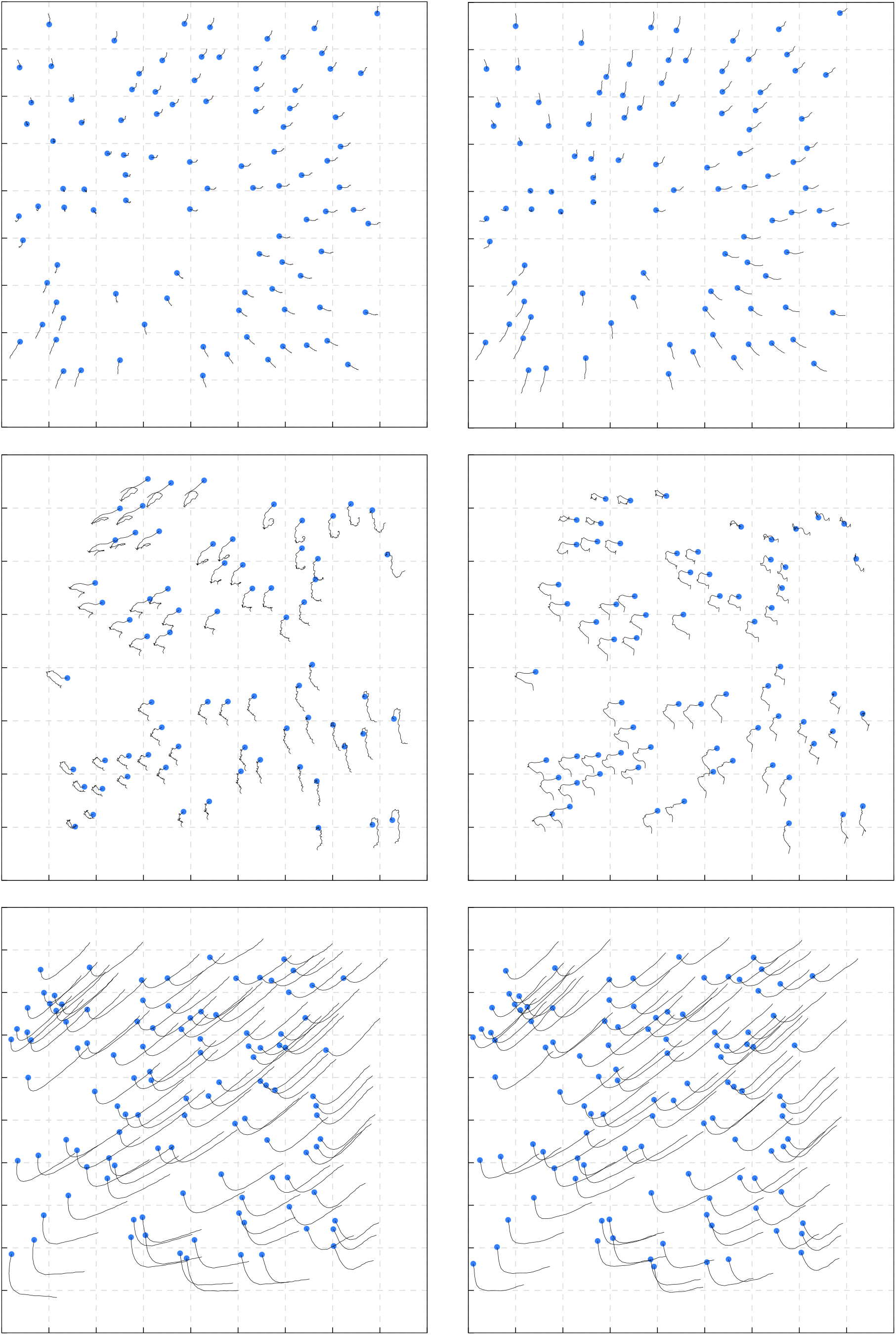
*Example trajectories using our own version of* MotionCor2 *(left) and Bayesian polishing (right) for the ribosome (top), *β*-galactosidase (center) and γ-secretase (bottom). Particle motion is scaled by a factor of* 30. *The blue dot indicates the start of the trajectory*.

### 3.3 B-factors

From the particle trajectories estimated by both Bayesian polishing and our implementation of *MotionCor2*, we computed the FCCs as defined in Eq. 20, and from those the *B_f_*, *C_f_* and *D_κ_* factors. Since the three sums in Eq. 20 are updated after the alignment of each micrograph, once all of them have been aligned, the computation of the B-factors only takes fractions of a second. In the previous particle polishing implementation, this step would take up multiple days of additional CPU time to calculate two half-set reconstructions for each movie frame. A comparison between the B-factors obtained by the two methods are shown in Fig. 4. A comparison to the previously published B-factors is shown on the left-hand side of Fig. 6.

**Figure 4:**
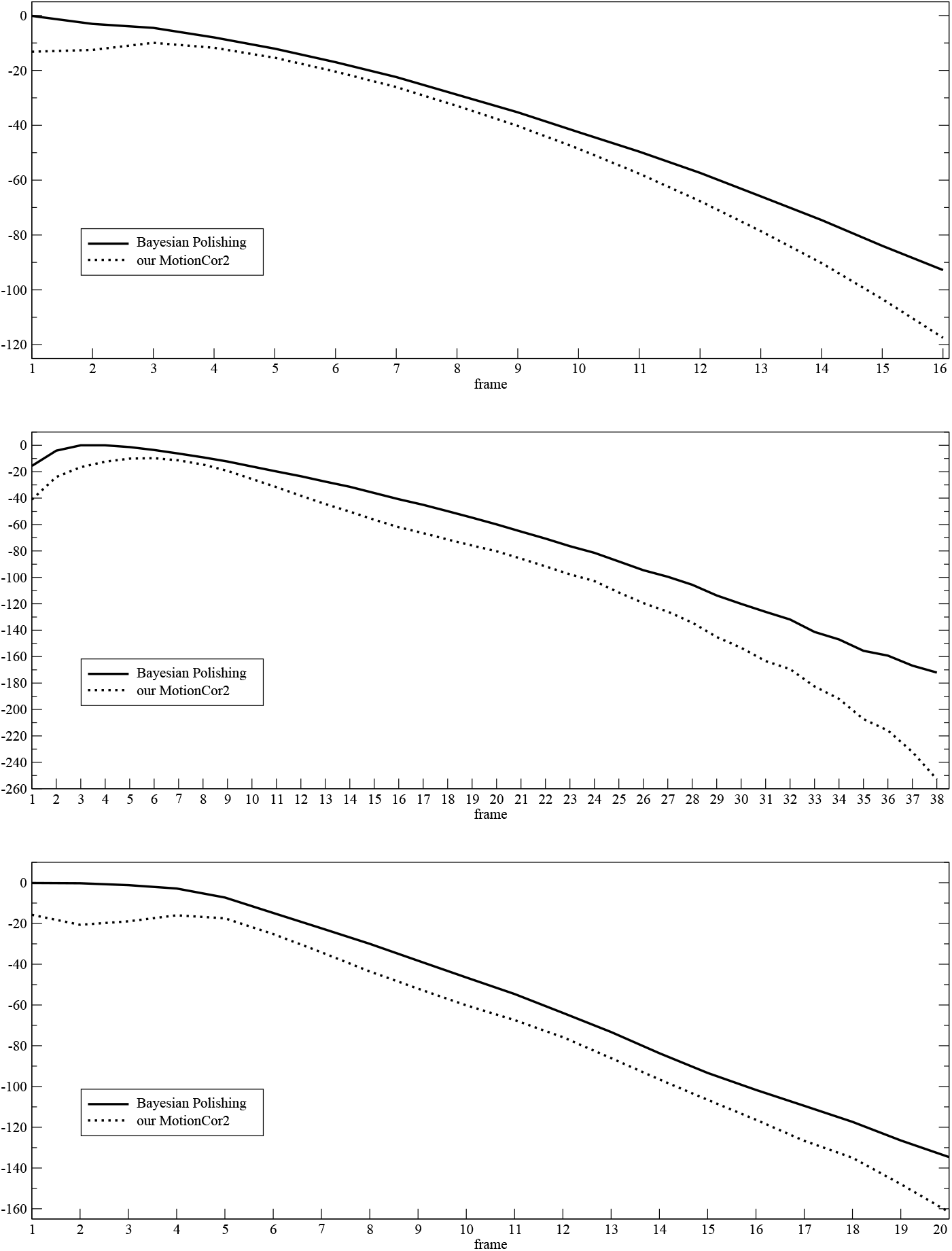
*Relative B-factors for the ribosome (top), β-galactosidase (center) and γ-secretase (bottom). The two sets of B-factors share the same D_κ_ factors, making their relative vertical position meaningful. The observation that the B-factors from the Bayesian polishing are higher than the ones from our* MotionCor2 *implementation suggest that Bayesian polishing models the motion more accurately*.

Generally, a set of relative B-factors can be shifted by a constant offset without altering the resulting pixel-weights. Such a shift corresponds to multiplying the *D_κ_* factors by a Gaussian over *κ*, and it cancels out when the division in Eq. 22 is performed. In order to make a meaningful comparison between the B-factors for motion estimated by Bayesian polishing and *MotionCor2*, we have estimated both sets of B-factor with the same *D_κ_* factors. This is equivalent to treating the movie frames aligned using Bayesian-polishing and those aligned using our implementation of the *MotionCor2* algorithm as a movie of twice the length. As can be seen in Fig. 4, the B-factors from Bayesian polishing are better over all frames for all three cases. The average improvement in B-factor over all frames is 9 Å^-2^ for the ribosome, 26 Å^-2^ for *β*-galactosidase, and 15 Å^-2^ for γ-secretase. These increases suggest that more high-resolution signal is present, and hence that Bayesian polishing models motion more accurately than the *MotionCor2* algorithm. We will confirm this in the following section.

To confirm that our new technique of estimating B-factors does not yield systematically different B-factors from the original method [21], we also calculated the B-factors using the original method, but with the trajectories from Bayesian polishing, for comparison. These plots are shown on the left-hand side of Fig. 6 and they indicate that the new technique produces values that are close to those obtained through the old one. The similarity between the two curves is especially striking for the ribosome dataset (top left in Fig. 6), where image contrast is the strongest. The greater smoothness of the curve obtained through the new technique in the *β*-galactosidase plot (center left in Fig. 6) indicates that the new technique is more robust than the old one. This is to be expected, since the linear Guinier-fit applied by the old technique [21] has to rely on the frequency range where the FSC is sufficiently large, and that range can become very small in later frames.

### 3.4 Resolution

Finally, the gold-standard FSCs are compared to those from the two *MotionCor2* implementations in Fig. 5 and to the previously published results on the right-hand side of Fig. 6.

**Figure 5:**
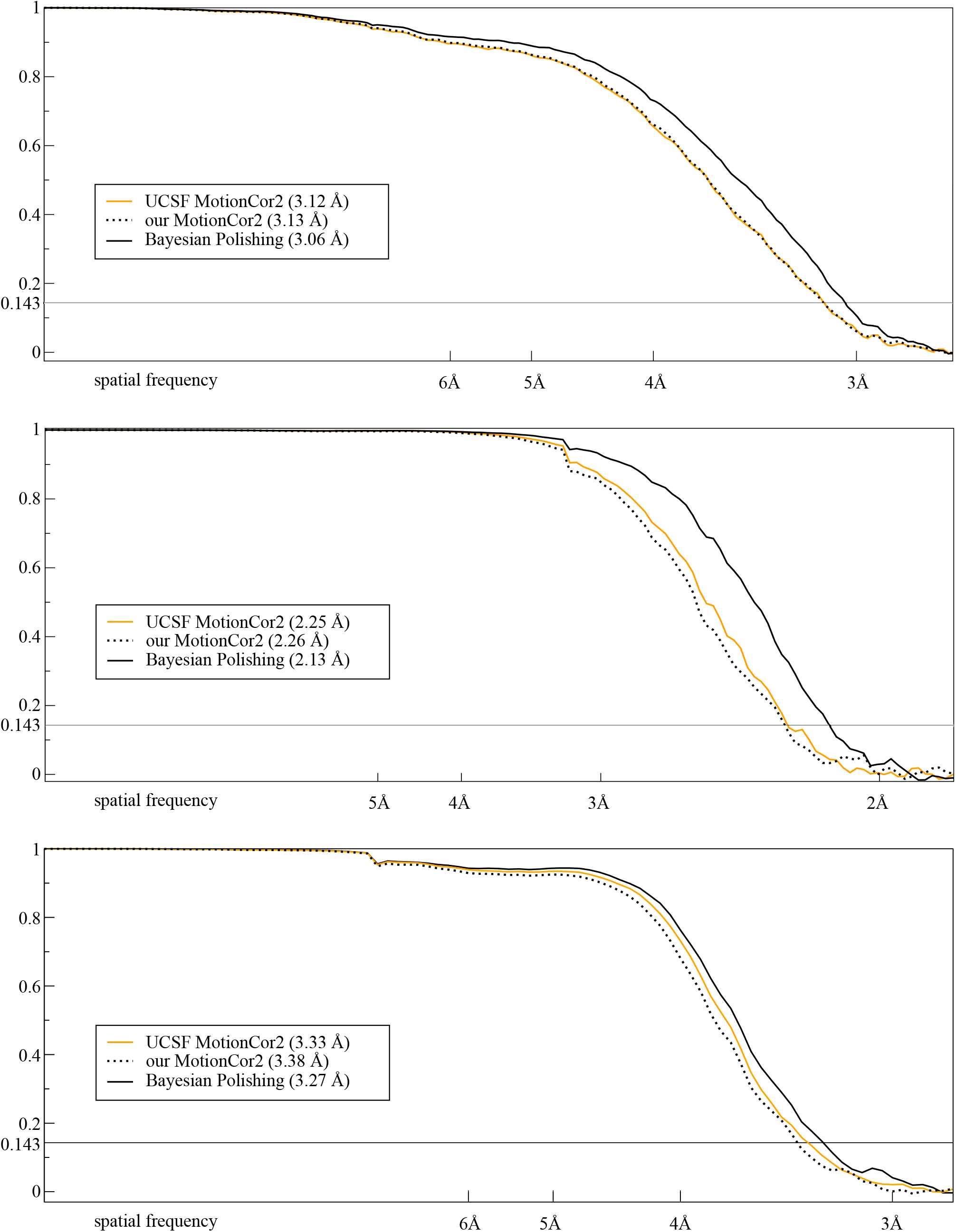
*Gold-standard FSC plots for the ribosome (top), β-galactosidase (center) and γ-secretase (bottom). The values in parentheses indicate the* 0.143-*FSC resolution. The continuous orange line results from the official UCSF implementation of* MotionCor2, *the dotted black line from our own*.

**Figure 6:**
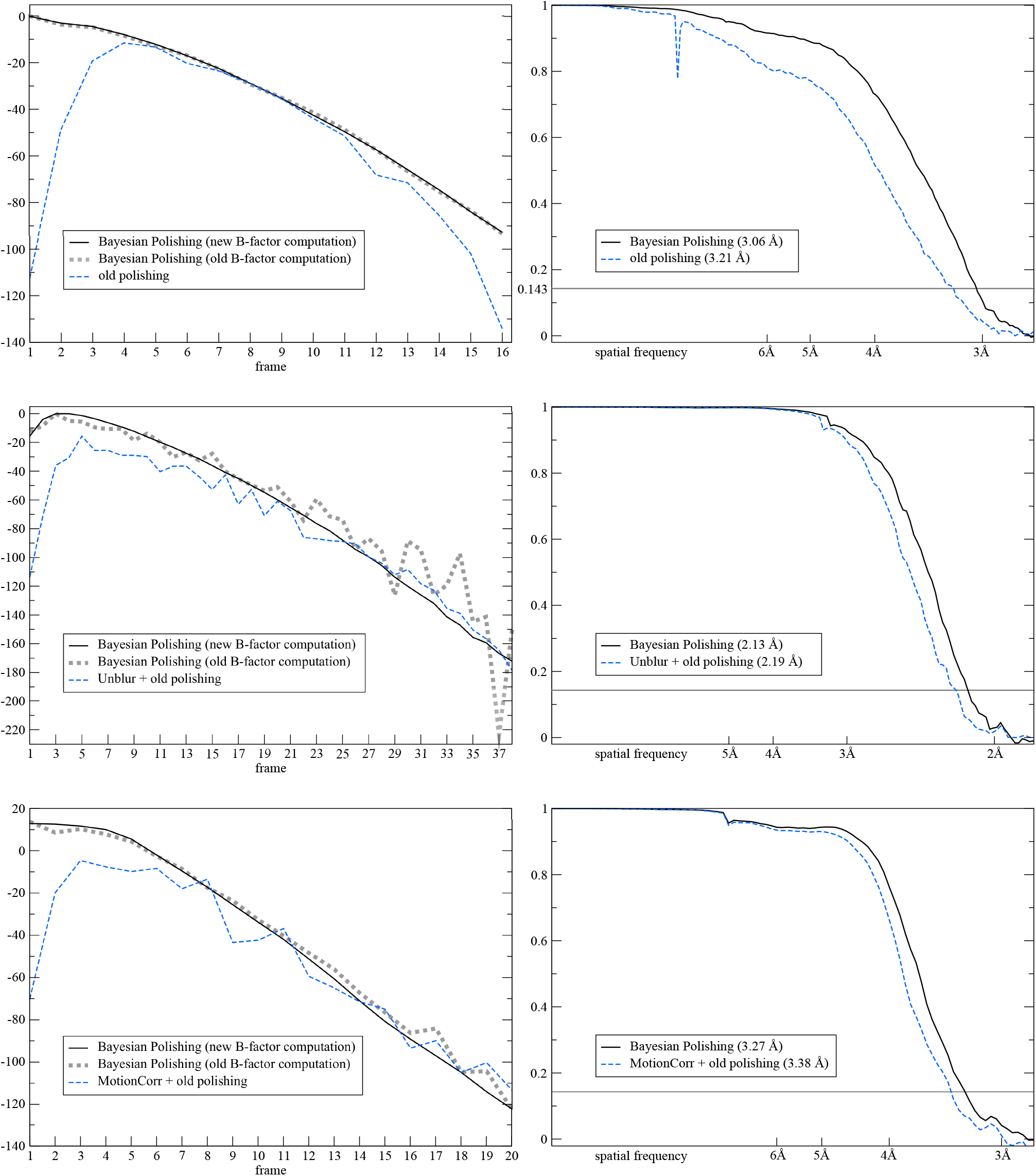
*Comparison to previously published results for the ribosome (top), β-galactosidase (center) and γ-secretase (bottom)*. **Left***: Relative B-factors. Unlike in Fig. 4, the vertical positions of these curves are arbitrary – only their shape holds any meaning. The continuous black and the dotted grey lines correspond to the same motion estimate, but they have been determined using the new and the old B-factor estimation technique, respectively. Their similarity indicates that the new technique estimates the same B-factors as the old one, albeit in a more robust way. The dashed blue line corresponds to previously published B-factors. Note the stark improvement in the beginning of the sequence*. **Right***: FSC curves comparing the new results to the previously published ones. Please note that the old polishing approach estimated the motion as superimposed over that estimated by another, reference-free method, while Bayesian polishing always works on the raw unaligned micrographs and aims to model the entire motion by itself*.

The FSCs were measured under the same solvent mask as had been used in the three previous publications and the effects of mask-induced correlation were corrected for through phase-randomisation [6] using RELION’s post-processing program. To further improve their precision, the resolutions indicated in the figures have been measured as the resolutions at which the linearly interpolated FSCs cross the 0.143 threshold.

As can be seen in the FSC plots, Bayesian polishing leads to an increase in resolution over both *MotionCor2* and the previously published results in all three cases. The increase over *MotionCor2* is the greatest for the *β*-galactosidase dataset. We assume that this is because that dataset extends to higher resolution than the other two data sets, and Bayesian polishing makes more efficient use of the high spatial frequencies by comparing the noisy movie frames with high-resolution reference projections. This assumption is further supported by the fact that *β*-galactosidase is also the only dataset on which traditional polishing applied after Unblur produces a better reconstruction than running *MotionCor2* alone. Compared to our previously published results, the increase in resolution is highest for the ribosome dataset. We assume that this is because of the high molecular weight of these particles, which allow for precise modelling of the motion tracks. The γ-secretase data set yields the smallest increases in resolution, both in comparison with *MotionCor2* and our previous results. Possible reasons for this will be discussed in the following.

We have also analysed the resolution of the resulting reconstructions as a function of the number of particles, as proposed by Rosenthal and Henderson [18]. These plots are shown for both our results and those obtained from the UCSF implementation of MotionCor2 in Fig. 7. They indicate that in order to reach the same maximum resolution with Bayesian Polishing as with the UCSF implementation of MotionCor2, only 66% of the particles are needed for the ribosome and only as little as 30% for *β*-galactosidase. For γ-secretase, only 60% of the particles are needed to reach the same *intermediate* resolutions, although the same number of particles are required to obtain the maximum resolution. This suggests that at high resolutions, γ-secretase is limited by additional effects beyond the experimental noise and the uncertainty in particle alignment. Such effects could include molecular heterogeneity, anisotropic magnification, an insufficient particle box size or variations in microscope parameters across the dataset. The latter is especially likely since this dataset was collected in six different sessions over a time span of half a year.

**Figure 7:**
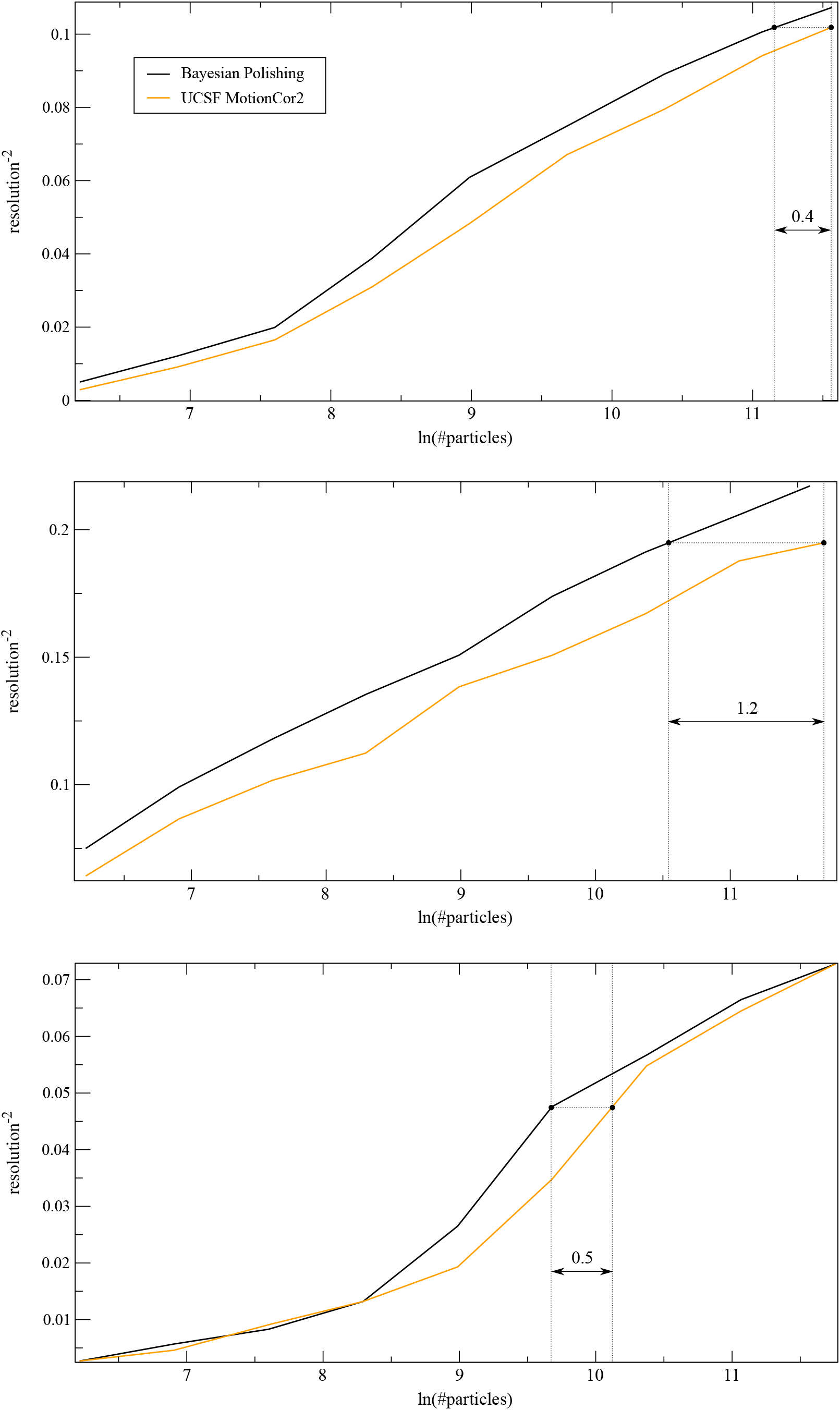
*Plot of the inverse-squared resolution as a function of the number of particles, as proposed by Rosenthal and Henderson [18] for the ribosome (top), β-galactosidase (center) and γ-secretase (bottom). The horizontal distance between the curves describes the fraction of the number of particles required to obtain the same resolution with Bayesian Polishing as with the UCSF implementation of MotionCor2. The indicated, distances correspond to 66%, 30% and 60% of the particles, respectively. Note that the horizontal distance shrinks to zero at the right-hand edge of the γ-secretase plot. This implies that the γ-secretase dataset is limited by additional effects at high resolutions*.

## 4 Conclusions

We have presented Bayesian polishing, a new method for the estimation of particle motion and of the corresponding per-frame relative B-factors. We have compared our method to *MotionCor2* and to the previously existing particle polishing method in RELION on three publicly available datasets. In all three cases, Bayesian polishing led to an increase in resolution over both alternatives. Since the FSC-based resolution estimates are influenced by many other factors besides particle motion, the accuracy of motion estimation was also measured by comparing the estimated relative B-factors. We have shown that Bayesian polishing produces better B-factors than our implementation of *MotionCor2* for all frames of all datasets, with an average improvement over all three data sets of 16 ^Å^A^2^, while the achieved resolution after refinement shows that our implementation of *MotionCor2* is comparable to the official UCSF implementation. The comparison of the shapes of our new B-factor curves to our previously published curves suggests that Bayesian Polishing captures significantly more of the initial motion than the existing particle polishing method in RELION. This allows the use of almost as much high-resolution data from the first few movie frames as from the intermediate movie frames, thereby obviating the need for the practice of discarding the first few movie frames [10]. Finally, we have shown that the new FCC-based technique of estimating B-factors measures the same B-factors as in the existing particle polishing method, but much faster and more robustly.

We have also presented a method that enables the user to determine the optimal parameters governing the statistics of motion. Since the final resolution of the resulting reconstructions appears to be relatively insensitive to those parameters, and the parameter hyper-optimisation algorithm requires considerable amounts of memory, we do not necessarily recommend estimating new parameters for each dataset. Instead, we expect that use of the default values should produce similar results, unless the dataset has been collected under unusual conditions. For example, re-estimating the motion parameters may be necessary for datasets that exhibit a much smaller fractional electron dose or significantly thinner or thicker ice, or if special grids are used that are designed to minimise beam-induced motion.

Bayesian polishing has been implemented as part of the open-source release of relion-3.0. The new implementation no longer requires storing aligned micrograph movies or movie particles, and is capable of performing on-the-fly gain correction on movies stored in compressed TIFF format. Thereby, the new implementation strongly reduces the storage requirements of performing particle polishing in relion. Because the new method has outperformed the previously existing particle polishing in all tests performed, the new approach replaces the old one on the graphical user interface (GUI) of relion-3.0.

## 5 Acknowledgements

We thank Jake Grimmett and Toby Darling for assistance with high-performance computing, and Christopher Russo and Richard Henderson for critical reading of the manuscript. This work was funded by the UK Medical Research Council (MC_UP_A025_1013 to SHWS) and the Swiss National Science Foundation (SNF: P2BSP2_168735 to JZ).

